# Computational Prediction of *Plasmodium falciparum* Antigen-T-cell Receptor Interactions via Molecular Docking: Implications for Malaria Vaccine Design

**DOI:** 10.64898/2026.03.18.712575

**Authors:** Gilbert Kipkoech, Wesley Kanda, Beatrice Irungu, Mary Nyangi, Cecilia Kimani, Ruth Nyangacha, Lucia Keter, Diana Atieno, Jeremiah Gathirwa, Elizabeth Kigondu, Edwin Murungi

## Abstract

Malaria is one of the deadliest diseases in sub-Saharan Africa and Southeast Asia. The majority of the fatalities occur mostly in children under 5 years and pregnant women and this is due to infection by *Plasmodium* spp, of which *Plasmodium falciparum* is the most virulent and is responsible for most of the morbidity and mortality. Despite various public health interventions such as use of insecticide-treated bed nets, spraying of homes with insecticides and use of WHO recommended artemisinin-based combination therapies (ACT), malaria prevention still faces major setback due to drug and insecticide resistance by *P. falciparum* and mosquitoes respectively. The study uses molecular docking and immunoinformatics to screen various *Plasmodium* spp antigens and evaluate their antigenicity and suitability as vaccine candidates. The *P. falciparum* antigens and T-cell receptor (TCR) structures were obtained from Protein Data Bank (PDB) based on a range of factors related to their role in the lifecycle of the parasite and their status as vaccine targets. Protein structures not available in the PDB were predicted using AlphaFold. The 3D structures of selected *P. falciparum* antigens and TCR structures were downloaded in PDB format then all water molecules, Hetatm, and bound ligands were deleted from the protein structures using BIOVIA Discovery Studio Visualizer. Subsequently, molecular docking was done using ClusPro v2.0 server and docked complexes were compared. The findings of this study gave valuable insights into the interaction of human immune response with *P. falciparum* antigens. The best three ranked antigen complexes are PfCyRPA, PfMSP10 and PfCSP and this confirm their use as potential candidates for vaccine development. This study highlights the usefulness of computational docking in identifying *P. falciparum* antigens of excellent immunogenic potential as vaccine candidates.

## Introduction

### Background on malaria and Plasmodium falciparum

The global burden of malaria remains huge particularly in sub-Saharan Africa and Southeast Asia where majority of the fatalities occur mostly in children under 5 years and pregnant women [1]. According to World Health Organization (WHO) report of 2022, it is estimated that there were 249 million cases of malaria with 608,000 deaths reported in 85 countries. African nations are immensely disproportionately affected and account for 94% of the total reported cases and 95% of deaths [2]. The causative agent for malaria is *Plasmodium* spp, of which *Plasmodium falciparum* is the most virulent and is responsible for most of the morbidity and mortality. Infection transmission is initiated when infected female anopheles mosquitoes bite non-infected humans transferring sporozoites that undergo developmental stages in humans to become infective. *Anopheles gambiae* is the dominant transmission vector in most areas [3]. Despite an array of public health interventions aimed at halting transmission such as insecticide-treated bed nets, spraying of homes with insecticides and use of WHO recommended artemisinin-based combination therapies (ACT), malaria prevention faces the main setback drug and insecticide resistance by *P. falciparum* and mosquitoes respectively. Thus, the development of novel malaria interventions is a pressing priority.

The life cycle of *Plasmodium falciparum* is distinctly complex, comprising of numerous stages in the mosquito and human hosts [4]. When entering the human host, *P. falciparum* infects red blood cells, which are killed and thus start causing symptoms of severe disease, including anemia, cerebral malaria, and multi-organ failure [5,6]. It is worth noting that besides being resistant to majority of the traditional antimalarial agents, the pathogen is also able to evade human immunity by antigenic variation [7]. In addition to that, *P. falciparum* exhibits extreme genetic diversity that allows it to adjust to various environments via population processes [8].

### Importance of Vaccine Development

Significant progress has been made in creating malaria vaccines, most notably the RTS,S/AS01 vaccine [9] that has been reported to decrease cases of clinical malaria (by approximately 56 to 60 per cent) and severe malaria (by approximately 47 to 60 per cent) in children aged 5-17 years one year after vaccination [10]. Moreover, by providing herd immunity, the RTS,S//AS01 vaccine can also diminish malaria transmission within populations [11]. Mathematical modeling estimates that were the vaccine’s coverage to reach the levels attained during routine vaccination of children, immense reduction in malaria deaths would be witnessed [12]. Such efficacy points to the possibility of halting malaria through immunisation as a complementary measure to transmission control and treatment [13].

Despite the encouraging trend, several outstanding issues remain to be resolved in the search for a universally efficacious malaria vaccine among them the cost of development [13]. Furthermore, public awareness of, and acceptance of the vaccine is also critical in the ultimate successful roll-out. Communities within malaria-endemic districts have been reported to have a positive view of immunization, with most caregivers responding that they will immunize children against malaria [14,15]. But misinformation and poor awareness can deter acceptance, making transparent measures critical to teach people on the benefits of the vaccine [15,16].

Moreover, the RTS,S vaccine also has its effect on herd protection, where it can contribute to herd immunity and reduce malaria transmission within populations [11]. Mathematical modeling estimates that where coverage of the vaccine is reached up to levels that have been achieved by routine vaccinations of children, there can be significant reductions of malaria deaths and malaria cases [12]. Such possibilities reinforce the means by which malaria immunization can be a key component of overall population interventions for malaria prevention and, ultimately, malaria elimination.

### Role of T-Cell Receptors in Immune Response

T-cell receptors (TCRs) are of critical significance in malaria immunity, even more on *P. falciparum* infection. The role of TCR is to recognize the antigens in the major histocompatibility complex, and hence induce cell activation thus resulting in immune responses [17]. The activation of CD4+ and CD8+ cell subsets has a critical role in malaria defense, where cells coordinate cellular and humoral immune responses capable of containing and clearing the parasite [18].

The CD4+ T cells, also referred to as helper T cells, are also key in the coordination of the immune response. These cells engage in activating B cells, generating antibodies, and activating the cytotoxic action of the CD8+ T cells, directly killing infested cells [19]. During malaria infection, the CD4+ T cells can generate multi-lineages of cells, such as the Th1 cells, of major importance for the control of malaria infection with Plasmodium via generation of pro-inflammatory cytokines such as IFN-γ and TNF-α [20]. The action of the Th1 cells has been associated with parasitemia control and malaria infection outcomes. There also exists the Tr1 cells, a key subset of immunosuppressive CD4+ T cells, whose action inhibits protection against the parasite by the action of the Th1 cells, and enhances generation of infection and inhibits immunological disease of malaria infection [21].

Moreover, the dynamics of T-cells’ response to malaria are also influenced by infection chronicity. Repeated antigenic malaria stimuli can lead to exhaustion of B-cells and also of T-cells. It is demonstrated that frequent *P. falciparum* parasites exposure is followed by elevated levels of CD4 T-cells that exhibit phenotypic markers of exhaustiveness. It is evident on programmed cell death-1 (PD-1) alone and also co-expression of PD-1 along with lymphocyte-activation gene-3 (LAG-3). The proliferation of PD-1 and co-expression of PD-1/LAG-3 is of specific interest to CD45RA+ CD4 T-cells [22]. It is evident on animal models and also on humans where exhaustively differentiated T cells exhibit reduced cytokine generation along with proliferative activity [22]. It is important to understand these processes to determine means of enhancing responses of T-cells, e.g., by employing therapy that inhibits inhibitory pathways to restore function of T-cells.

In addition to CD4+ cells, CD8+ cells also make important contributions to malaria immunity. The cytotoxic cells can identify infected hepatocytes and red blood cells and destroy them, thereby inhibiting proliferation of the parasite [23,24]. It has been shown that malaria can be recognized by CD8+ cells through cross-presentation, enhancing their ability to respond to infection of the blood stage [25]. The activity of the CD8+ cells also relies on memory characteristics, and these can be changed with history of infection, and with access of specific antigen [26].

### Molecular Docking and Immunoinformatics

Molecular docking and immunoinformatics have been critical drug discovery tools. The technologies have contributed immensely in understanding how medicines combat some of the epidemic diseases such as COVID-19 and malaria. Molecular docking is a computer program which first made its entry in the 1980s in an effort of making it possible for addressing best pose of a target molecule when complexing a receptor [27]. The first program for molecular docking, however, was developed by Kuntz and his group in 1980s [28]. Molecular docking revolutionized understanding of functions of a molecule on a molecular scale [27]. It made it possible for complexation of a molecule and a receptor, calculation of affinities of complexation, and actual medicine designing [29,30]. It accelerates drug discovery making it possible for screening of thousands of compounds for identifying a potential medicine and reducing effort and time in confirming results in experiments [31].

On the other hand, immunoinformatics, or computational immunology, employs computational strategies to analyze and predict immune reactions with a major focus on the design of vaccines. Immunoinformatics has recently emerged as an important tool in immunological studies, which include vaccine design and development. Despite that this technology is still evolving, the computational models have played a significant role during the selection of antigens or proteins and complex immunologic data analysis, thus facilitating the formulation of new testable hypotheses [32]. Immunoinformatics is established on the knowledge of the antigens’ epitopes that are the targets of the immune response to develop multi-epitopic vaccines that can elicit strong immune reactions [33]. One of the key areas of immunoinformatic applications is the prediction of B-and T-cell epitopes by computational strategies to aid the development of vaccines that can activate adaptive immunity [34]. It is particularly relevant to emerging pathogens like the SARS-CoV-2 where classical strategies to the development of vaccines can fail [35]. The combination of immunoinformatic analysis with molecular docking enables the rational peptide vaccine designing by predicting the affinities of the peptide-MHC interaction, their recognition by the major histocompatibility complex (MHC) molecules, and by the T cells. These techniques allow for selection of best conformations that can be validated through experimental studies. The molecular docking also enables screening of the potential vaccines to interact with the Toll-like receptors (TLRs), a major component of initiating the innate immunity [35]. Numerous studies have reported that reformulating the key proteins used in the vaccine improves efficacy and immunogenicity thus improving response and protection.

Computational biology has gained popularity in the recent past and it has accelerated drug discovery and development. The latest technologies involving artificial intelligence (AI) and deep learning have been widely used on research platforms, thus accelerating drug discovery. Deep learning is based on the idea of artificial neural networks (ANNs) that can resemble the brain learning process [36]. In most occasions, AI, that assists in predicting drug-targets, has significantly enhanced drug discovery by molecular docking and immunology. In this way, over a single conformation is obtained and the best of them are chosen depending on their binding capacity. The strategy has helped in developing vaccines and drugs significantly.

### Objective of the study

This research was conducted with the purpose of applying molecular docking and immunoinformatics to identify the most appropriate drug-target orientations. The study explicitly takes into account the interaction of the antigen *P. falciparum* with the T-cell receptor in order to identify the majority of the most appropriate malaria vaccine targets. Following the molecular docking, binding affinities and interfaces of the complexes were determined in an effort to determine the activation of immune responses. Antigen-receptor docks were ranked and the most active complexes, according to binding affinities, were chosen to undergo further investigations. The results obtained in this study give a starting point of experimental testing and vaccine development in malaria.

## Methodology

### Selection of Antigens

#### Criteria for Choosing Plasmodium falciparum Antigens

The *P. falciparum* antigens were chosen based on a range of factors related to their role in the lifecycle of the parasite and their status as vaccine targets. To achieve maximum coverage against malaria, for example, priority was given to antigens with widespread sequence conservation between different *P. falciparum* strains. Also, our study prioritized antigens with key roles in important biological processes, such as host cell invasion, host cell egression, and immune evasion. The reason for this prioritization was to maximize the probability of effective immune responses. Also, our study focused on secreted and surface-expressed antigens. Surface-expressed and secreted antigens are more likely targets for host immune systems. Based on previous studies, this study also considered the immunogenic potential of the antigens. Antigens that have shown promise in previous immunological studies (Table 1), eliciting strong immune responses in humans or model organisms, were considered.

**Table 1:** Summary of previous experimental studies validating Plasmodium falciparum antigens as vaccine candidates.

| Antigen | Key experimental study | Experimental approach | Main outcome |
| --- | --- | --- | --- |
| PfCSP | Mahmoudi et al., 2017 [37] | Phase III clinical trial in African children | Demonstrated relatively little efficacy towards malaria control |
| PfAARP | Wickramarachchi et al., 2008 [38] | Immunological characterization, sera from endemic area & invasion-inhibition assays | PfAARP-N reactive with immune sera and rabbit anti-AARP inhibited merozoite invasion. |
| PfRh5 | Reddy et al., 2014 [39] | Recombinant RH5 production and erythrocyte binding assays | RH5 binds erythrocytes similar to native protein - supports vaccine candidacy. |
| PfRipr | Takashima et al., 2022 [40] | PfRipr fragment vaccine, immunogenicity, and inhibitory antibodies | PfRipr5 fragment induced potent invasion-inhibitory antibodies against *P. falciparum* blood stage. |
| PfCyRPA | Williams et al., 2024 [41] | Blood-stage vaccine component study with CyRPA, RH5, RIPR | Demonstrated that CyRPA (with RH5/RIPR) is a conserved target and elicits inhibitory antibodies. |
| PfMSP10 | Bendezu et al., 2019 [42] | Serological assays in malaria-exposed individuals | MSP10 found to be immunoreactive; supports its potential as a vaccine or serological marker. |
| PfSEA-1 | Raj et al., 2014 [43] | Animal immunization and in vitro egress assays | Antibodies to PfSEA-1 reduce parasite replication by arresting schizont rupture. |

#### Sources of Antigen Sequences

The 3D structures of *P. falciparum* antigens were obtained from the Protein Data Bank (PDB) (https://www.rcsb.org/). The PDB is a database with structures of proteins that are experimentally determined using different techniques such as through X-ray crystallography or cryo-electron microscopy. However, some of the protein structures are not available the RCSB Protein Data Bank and therefore their computational models were predicted using AlphaFold Structure Database (https://alphafold.ebi.ac.uk/). There are numerous antigen candidates that can be explored for malaria vaccine development. Table 1 consists of some of the potential candidate target proteins that can be modelled for vaccine development.

### T-cell Receptor and MHC Molecules Data

#### Criteria for Selection T-cell Receptor and MHC molecules

In this study, only TCRs known to interact with malaria-specific antigens or closely related epitopes were selected to ensure relevance to the *P. falciparum* infection. To simulate human immune responses, only TCR sequences from humans were chosen, as these receptors are crucial for understanding potential vaccine efficacy and T-cell mediated immunity in humans. In addition, preference was given to TCRs with resolved crystal structures or high-quality modeled structures to enable accurate docking simulations.

When choosing between TCR1 and TCR2 for docking simulation, it is important to consider their roles and prevalence. TCR1 (Gamma-Delta TCR) is less common type, found in about 1-5% of T cells, and is primarily located in mucosal tissues and skin. It has a function in localized immune reactions and is still not fully understood. TCR2 (Alpha-Beta TCR) is found in over 90% of T cells and participates in the majority of immune responses. It is the conventional TCR that interacts with peptide-MHC complexes to activate T cells [44]. For most studies, TCR2 (Alpha-Beta TCR) is the preferred choice due to its prevalence and well-characterized role in immune responses.

MHC Class II molecules were selected in this study. For malaria vaccine design, MHC Class II molecules are often studied. They stimulate helper T-cell and antibody-mediated immunity thus they play a crucial role in generating a strong and sustained immune response, which is vital for long-term protection against the parasite.

#### Database and Sources for T-cell Receptor Structure and MHC molecules

The T-cell receptor (TCR) sequences and structures were retrieved from the Protein Data Bank (PDB) (https://www.rcsb.org/). The PDB was specifically used to obtain experimentally determined structures of human TCRs bound to various antigen-MHC complexes, particularly those involved in recognizing malaria epitopes. In this study, TCR2 (Alpha-Beta TCR) (PDB ID: 4WW1) was selected. MHC Class II was source from PDB (PDB ID: 3L6F)

### Preparation of Antigen and T-cell Receptor Structures

Proper preparation of protein structures is critical to the success of molecular docking studies. The 3D structures of selected *P. falciparum* antigens and TCR structures were downloaded in PDB format from the Protein Data Bank (PDB) and AlphaFold Structure Database. All water molecules, Hetatm, and bound ligands were deleted from the protein structures using BIOVIA Discovery Studio Visualizer v24.1.0.23298.

#### Molecular Docking Protocol

The Plasmodium antigens were docked to MHC Class II and further to TCR2 by means of ClusPro v2.0 server available on https://cluspro.org. It is a popular open source server for predicting diverse protein-protein interaction. It was selected because it provides a stable protein-protein docking server with a multi-step approach employing the use of the Fast Fourier Transform (FFT). Apart from that, the tool is capable of clustering the result based on binding energy and thus enable the determination of probable structures of the two protein bindings [45]. ClusPro is a web server whose basic home page is used for general purposes. Only two files are accepted by the server in the PDB format [46]. After the natural process of immune response, stepwise docking protocol was conducted in this research in two steps. The antigens of *P. falciparum*, in step one, were docked against Human MHC Class II protein. Once the initial docking was done, the binding energies and compatibility were determined using the docking energy scores. The server could form complexes of various affinities and cluster populace. The docking pattern is a notable procedure that is used in the prediction of affinities of the antigens to the MHC molecules preceding the presentation of the antigens to the T cells. This was formed with the assistance of natural immune mechanisms whereby it conjugates the antigens with MHC molecules which are then collected by the T cell receptors [47]. The most preferred MHC-antigen complexes that scored highly were selected following successful phase one docking and docking to TCR2.

In a bid to test the possible recognition and activation of T cells, the pre-programming of antigen-MHC complex with TCR2 was done. The significance of MHC molecules is extremely high during the presentation of processed plasmodium antigenic peptides to TCR2. This in turn triggers certain immune reactions. The protocol permitted examination of each antigen-MHC complex’s possibility for immunogenicity, simulating natural antigen recognition and presentation to T cells. Docking simulations for each were monitored through the ClusPro dashboard for any possible computational flaw.

### Data Analysis

Docking scores for all the complexes from ClusPro v2.0 were compared. Four energy parameter sets, namely, balanced, electrostatics favored, hydrophobic favored, and van der Waals + electrostatics, were used to execute the models after successful docking. The models were then ranked based on the cluster populations of the models, where a cluster’s population represented the probability of a given conformation. The model which belonged to the center of the biggest cluster of the hydrophobic-favored energy set was selected for analysis. The reason behind the selection of this model is that it has a large number of clusters and is biologically relevant as it is likely to give an accurate depiction of the protein-protein interaction. The structural and functional integrity of the predicted complex was checked by visualizing and validating the chosen model with the help of BIOVIA Discovery Studio Visualizer. The models that were ranked highest were further investigated to determine the essential interactions that occurred at the antigen-TCR interface including hydrogen bonds, salt bridges and van der Waals forces.

## Results

The docking of proteins-proteins was done through ClusPro web platform and 30 models were produced in each protein-protein complex. These models were classified into clusters according to structural similarity and the clustering data were examined to give the most reasonable binding conformations.

Table 3 presents the results of the clusters, cluster population, weighted score, and the minimum set of energy parameter. The Hydrophobic-favored parameter set of energy was prevalent in most of the models of interest, with the exception of the MHC-PfRh5-TCR apparently with the lowest energy model with the van der Waals + Electrostatics (VdW+Elec) parameter set. Cluster populations were diverse with the largest cluster size of MHC-PfCyRPA-TCR (174 members), MHC-PfMSP10-TCR (156 members) and MHC-PfCSP-TCR (154 members). It is interesting to note that MHC-PfSEA1-TCR possessed the lowest weighted energy score (-1185.3) indicating a very favorable binding conformation.

The model of choice took into consideration the cluster size more than the energy scores because bigger clusters have a better chance of reflecting stable binding configurations [46]. Cluster 0 from the Hydrophobic-favored Set was chosen for most complexes due to its high cluster population and relatively lower energy values. This aligns with previous observations that larger clusters correspond to energetically favorable binding landscapes. The PIPER energy Function for hydrophobic-favored and VdW+Elec are *E = 0*.*40E*_*rep*_ *+ − 0*.*40E*_*att*_ *+ 600E*_*elec*_ *+ 2*.*00E*_*DARS*_ and *E = 0*.*40E*_*rep*_ *+ − 0*.*10E*_*att*_ *+ 600E*_*elec*_ *+ 0*.*00E*_*DARS*_ respectively.

The selected protein-protein complexes were further analyzed by visualizing their three-dimensional structures using BIOVIA Discovery Studio Visualizer. Figure 1 below shows *P. falciparum* antigens in complex with MHC Class II and TCR2. Structural analysis of the selected complexes was done using PDBsum (https://www.ebi.ac.uk/thornton-srv/databases/pdbsum/). The best 3 ranked antigen complexes (PfCyRPA, PfMSP10 and PfCSP) were analyzed for key interactions, including hydrophobic contacts and hydrogen bonding at the interface. This helps in understanding the stability of the interaction.

**Figure 1.**
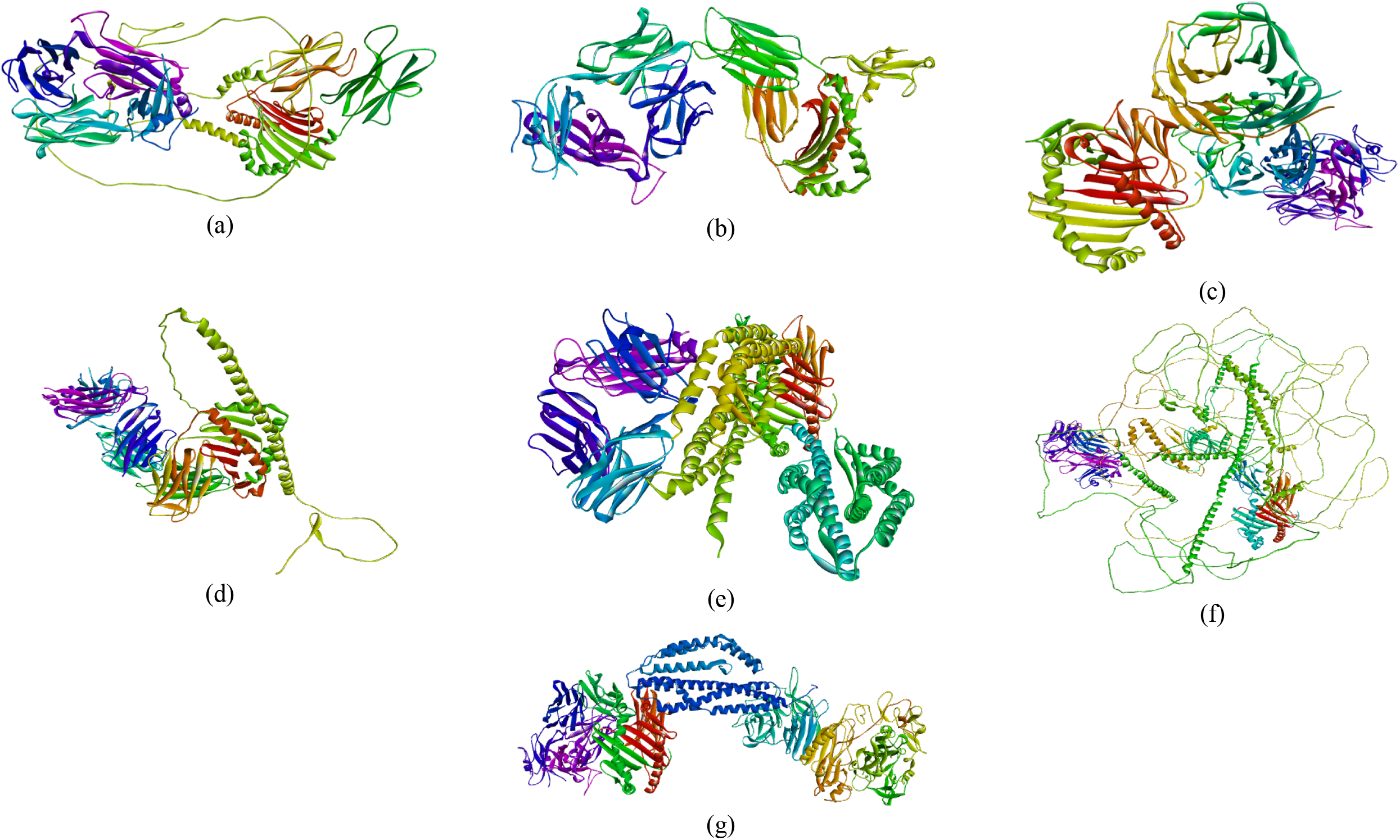
Structural representations of the best-docked conformations of Pf antigens-MHC-TCR complexes visualized using BIOVIA Discovery Studio. The models correspond to the cluster with high population and lowest-energy docked structures for Plasmodium antigens (a) PfAARP, (b) PfCSP, (c) CxRPA, (d) PfMSP10, (e) PfRh5, (f) PfSEA-1, and (g) PfRipr. The ribbon structures are color-coded to highlight secondary structural elements and interaction interfaces.

To assess the stereochemical quality and structural stability of the predicted complexes, Ramachandran plot analysis was performed (Figure 2). According to standard validation criteria, a high-quality model must have over 90% of residues in the most favored regions, on the basis of analysis of 118 structures with a resolution of at least 2.0 Å and an R-factor of not more than 20.0.

**Figure 2:**
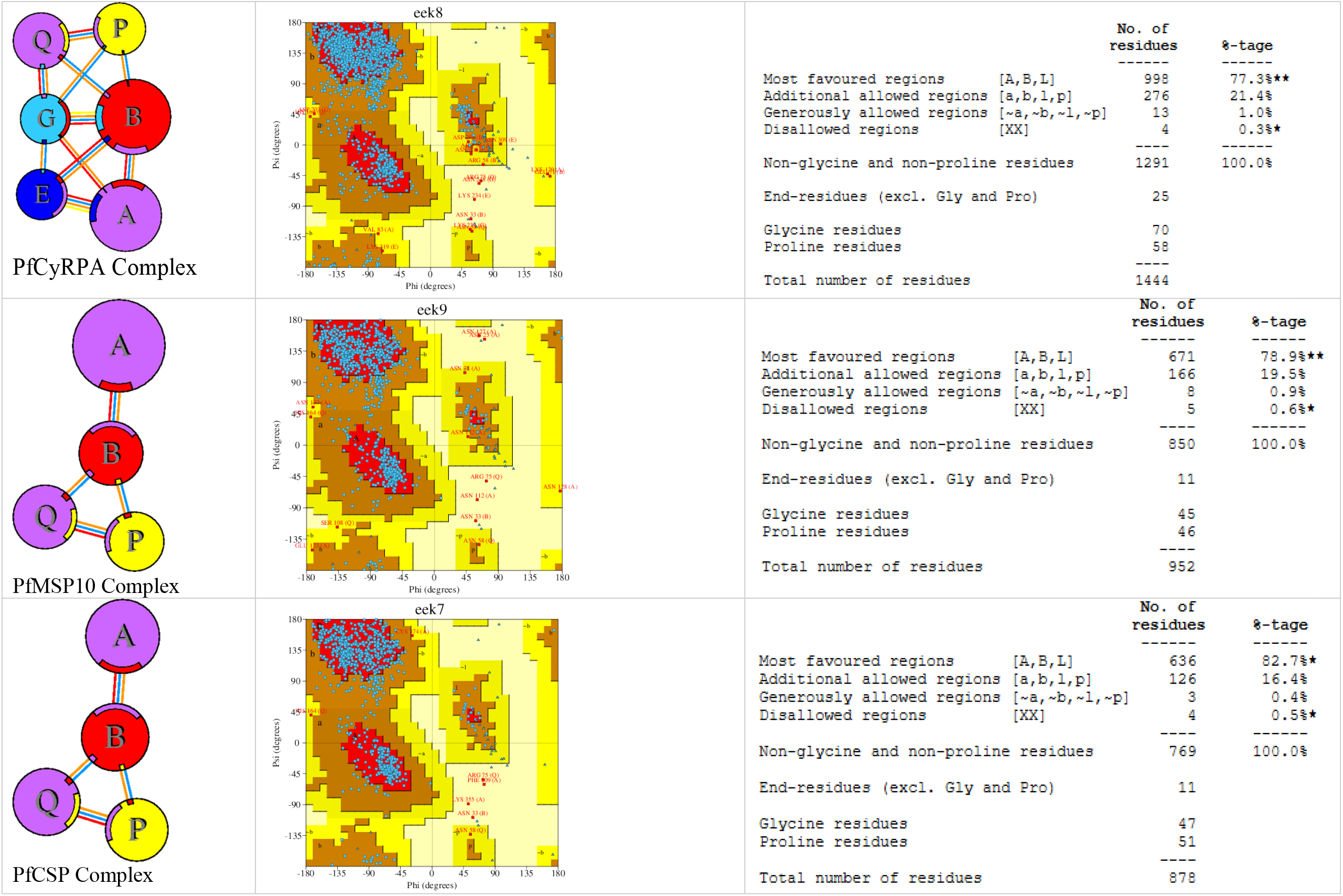
An illustration of how different protein chains interact (column 1). Chains interact with each other using different type of interaction as indicated by colored lines. Each circle’s size is proportional to the surface area of its corresponding protein chain. The area of the interface on each chain is shown as a colored wedge whose color is the same as the color of the other chain and whose size is proportional to the interface surface area. The Ramachandran plot (column 2) and statistics (column 3) for each complex shows the structural validation and overall stereochemical quality of the mod1e2led structures following protein-protein docking.

- **PfCSP Complex**: 82.7% of residues fall within the most favored regions, with 16.4% in additionally allowed regions and only 0.5% in disallowed regions.
- **PfMSP10 Complex**: 78.9% of residues are in the most favored regions, 19.5% in additionally allowed regions, and 0.6% in disallowed regions.
- **CyRPA Complex**: 77.3% of residues are within the most favored regions, 21.4% in additionally allowed regions, and 0.3% in disallowed regions.

While none of the models exceed the 90% threshold for the most favored regions, the presence of a high percentage in additionally allowed regions suggests that the structures remain well-refined and acceptable for further analysis. The low percentage of disallowed residues indicates minimal steric clashes, supporting the reliability of the predicted complexes.

Statistics for all the interfaces are given below (Table 2, 3 and 4). The interaction analysis of the top docked complexes reveals key stabilizing forces. In the PfCyRPA complex, non-bonded contacts dominate, with a striking 467 interactions in the primary interface. Similarly, PfCSP exhibits 439 non-bonded contacts, while PfMSP10 shows 464. Hydrogen bonds are also notable, with CyRPA forming 52 in its major interface, followed by PfMSP10 (46) and PfCSP (45). Among electrostatic interactions, PfCyRPA stands out with 13 salt bridges, the highest among the three complexes. These results highlight the importance of non-bonded contacts and hydrogen bonds in stabilizing the complexes, with salt bridges playing a crucial role in PfCyRPA.

**Table 2:** Some of the potential Plasmodium falciparum antigens considered for vaccine development.

| **Antigen** | **Source** | **Identifier** |
| --- | --- | --- |
| PfCSP (*P. falciparum* circumsporozoite protein) | PDB | 3VDJ |
| PfSEA-1 (Schizont Egress Antigen 1) | AlphaFold | AF-A0A143ZXM2-F1 |
| PfAARP (Apical Asparagine-Rich Protein) | AlphaFold | AF-Q8IFN2-F1 |
| PfRh5 (Reticulocyte Binding Protein Homolog 5) | PDB | 4U0Q |
| PfRipr (Rh5 Interacting Protein) | PDB | 8CDD |
| PfMSP10 (Merozoite Surface Protein 10) | AlphaFold | AF-J7GNL4-F1 |
| PfCyRPA (Cysteine-Rich Protective Antigen) | PDB | 5EZN |

**Table 3:** Summary of ClusPro Docking Results forAntigen-MHC-TCR Complexes.

| Complex | Cluster population | Weighted Score | Energy Parameter Set (Lowest Energy) |
| --- | --- | --- | --- |
| MHC-PfSEA-1-TCR | 104 | -1185.3 | Hydrophobic-favored |
| MHC-PfRipr-TCR | 112 | -834.7 | Hydrophobic-favored |
| MHC-PfRh5-TCR | 103 | -265.3 | VdW+Elec |
| MHC-PfMSP10-TCR | 156 | -806.7 | Hydrophobic-favored |
| MHC-PfCyRPA-TCR | 174 | -948.4 | Hydrophobic-favored |
| MHC-PfCSP-TCR | 154 | -806.5 | Hydrophobic-favored |
| MHC-PfAARP-TCR | 79 | -1037.9 | Hydrophobic-favored |

**Table 4:** Interface statistics for PfCyRPA complex.

| Chains | No. of interface residues | Interface area (Å<sup>2</sup>) | No. of salt bridges | No. of disulphide bonds | No. of hydrogen bonds | No. of non-bonded contacts |
| --- | --- | --- | --- | --- | --- | --- |
| A:H:B | 74:57 | 3148:3348 | 13 | - | 52 | 467 |
| A:H:E | 57:57 | 2693:2628 | 5 | 1 | 41 | 438 |
| A:H:G | 1:1 | 37:44 | 1 | - | - | 5 |
| B:H:E | 11:8 | 453:516 | 1 | - | 7 | 58 |
| B:H:G | 52:54 | 2526:2517 | 4 | 1 | 36 | 390 |
| B:H:P | 2:2 | 106:107 | - | - | 1 | 7 |
| B:H:Q | 6:8 | 380:326 | - | - | 1 | 38 |
| E:H:G | 5:6 | 246:275 | - | - | 3 | 41 |
| G:H:P | 3:2 | 114:128 | - | - | 2 | 12 |
| G:H:Q | 6:7 | 321:321 | 1 | - | 2 | 37 |
| P:H:Q | 52:50 | 2173:2166 | 6 | - | 32 | 385 |

**Table 5:** Interface statistics for PfCSP complex.

| Chains | No. of interface residues | Interface area (Å<sup>2</sup>) | No. of salt bridges | No. of disulphide bonds | No. of hydrogen bonds | No. of non-bonded contacts |
| --- | --- | --- | --- | --- | --- | --- |
| A:H:B | 66:50 | 2916:3069 | 7 | - | 45 | 439 |
| B:H:P | 7:5 | 288:323 | - | - | 3 | 43 |
| B:H:Q | 6:8 | 418:405 | - | - | 3 | 35 |
| P:H:Q | 51:55 | 2225:2175 | 6 | - | 31 | 406 |

**Table 6:**
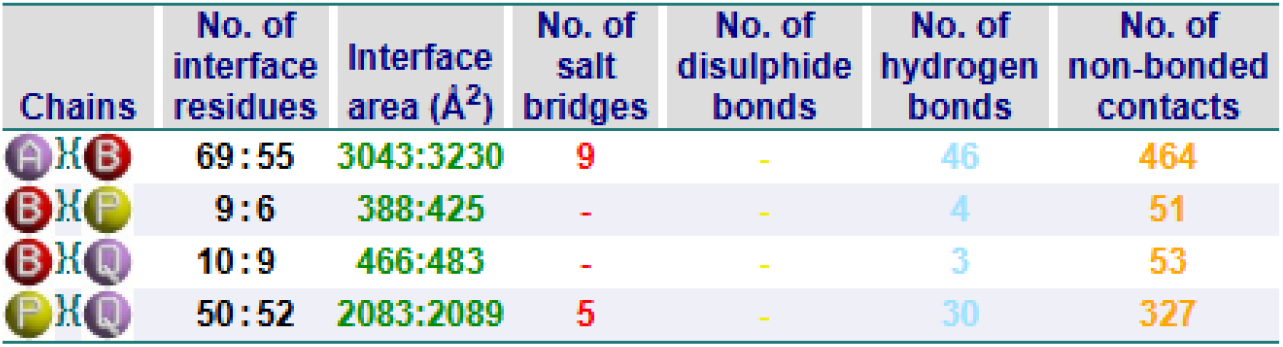
Interface statistics for PfMSP10 complex.

| Chains | No. of interface residues | Interface area (Å<sup>2</sup>) | No. of salt bridges | No. of disulphide bonds | No. of hydrogen bonds | No. of non-bonded contacts |
| --- | --- | --- | --- | --- | --- | --- |
| A:H:B | 69:55 | 3043:3230 | 9 | - | 46 | 464 |
| B:H:P | 9:6 | 388:425 | - | - | 4 | 51 |
| B:H:Q | 10:9 | 466:483 | - | - | 3 | 53 |
| P:H:Q | 50:52 | 2083:2089 | 5 | - | 30 | 327 |

## Discussion

Although the current study involves the application of computational docking to the study of antigen-receptor interactions, it is worth pointing out that some of the antigens used in the study are already known to be effective vaccine candidates that have been experimentally proven in prior studies. Table 1 is a summary of published articles which give immunological or functional support of the vaccine-relevance of each antigen. An example is PfCSP vaccines (RTS,S/AS01), which have demonstrated partial protection in children [48], PfAARP and PfRipr fragments produce invasion-inhibitory antibodies and the RH5 -CyRPA -RIPR complex (including PfCyRPA) is a conserved blood-stage target. The consistency of our computational predictions with this existing body of prior experimental evidence enhances the biological plausibility of the candidate antigens and gives a more explicit rationale in the validation of our hypothesis in subsequent in vivo and in vitro experiments.

A comparison of our docking results with published immunoinformatics studies was done to put our findings into perspective. As an example, immunoinformatics vaccine candidates docked to TLR4 can easily generate dozens of docked poses (clusters) with binding energies of the order of -1,000 to -1,500 kcal/mol. One study found that a malaria multi-epitope vaccine, which was docked to human TLR4, generated 30 clusters, the biggest cluster (cluster 0) containing 34 members and lowest-energy score of -1262.3 [49]. In another study, 30 models were generated and most optimal one contained energy -1514.2 (cluster center -1413.7) [50]. These values fall within the range of our values indicating that our docking energies are consistent with the previous studies. Kozakov et al. [46] also noted that larger clusters typically suggest more credible models.

The findings of this study give valuable insights into the interaction of human immune response with *P. falciparum* antigens, of utmost significance for vaccine research. The critical antigenic interactions were identified from the molecular docking simulations. These interactions can be of use for future malaria vaccine design. This study yielded a total of seven complexes with different cluster population.

One of the significant outputs of this research is the strong binding interaction of PfCSP with MHC molecule and T cell receptor. CSP is an important antigen that has been widely studied and has been well-characterized. It is promising and a key component of the novel RTS,S/AS01 malaria vaccine (Mosquirix). CSP is implicated at the beginning of *P. falciparum* infection to support the parasite’s invasion of the liver cells [51]. The high number of cluster population or members and the low energy scores obtained in this study enhances the biological value of CSP as a target in the vaccine design. Incorporation of the CSP into the RTS,S vaccine has been reported to offer protection against 56% of the case of the clinical malaria among 5 to 17-month-old infants [9]. The outcome of this research also warrants the continuation of the research to develop and further improve the effectiveness of the CSP-based vaccines.

The other key antigen that was studied is *P. falciparum* merozoite Rh5 interacting protein (PfRipr). This antigen mostly associates with Rh5 and CyRPA during the invasion of the erythrocyte. Docking results revealed promising interaction between MHC, PfRipr, and TCR. This signifies its potential to serve as a strong immunogen target. According to recent studies, PfRipr has been fully characterized and it is a nonredundant protein. This protein plays a critical role in red blood cell invasion. PfRipr is more conserved among proteins. These features make PfRipr an important candidate especially for the creation of blood-stage malaria vaccines [52].

Because PfRipr happens in a complex with Rh5 and CyRPA, the interactions need to be included within the molecular docking simulations to preserve the native binding conformation [53] and preserve the natural sequence of interaction. Exclusion of molecules can potentially change the structural stability of the binding site. This may impact the general efficacy of the candidate vaccine. The prospect of PfRipr as a future blood-stage vaccine target has already been shown by a number of studies including Ntege et al., [54], Williams [55], Correia et al. [56] and Takashima et al. [40]. Another key finding was noted with PfSEA-1. PfSEA-1 docking was found to give a small cluster with very low energy. A possible interpretation is that binding pose in PfSEA-1 is quite favorable yet distinct. Large cluster size is generally an indicator of confidence in ClusPro (when there are many similar low-energy poses) and a small cluster (even if low-energy) can indicate a lower count of repeated solutions. Kozakov et al. [46] note that cluster size is normally a more useful ranking measure compared to raw energy. In this way, a small cluster with low energy (such as PfSEA-1) cannot be considered as a large cluster.

PfMSP10 is a significant protein located in the merozoite surface and apical end. It is a structurally made up of two epidermal growth factors (EGF) where its C-terminal is attached to the red blood cell membrane. The role of PfMSP10 is still not known, but it is believed to contribute to growth stimulation and protein – protein interaction in merozoites and gametocytes [42]. The comparison of MSP1 with MSP10 in this study also puts into perspective the aspect of variability of the antigens to be remembered while developing vaccines. Although MSP1 is a well-studied protein with a well-characterized target of the immune response, the extensive variability of the protein among *P. falciparum* parasites is a limitation to the potential as a vaccine candidate [57]. Nevertheless, MSP10 was revealed to have strong interaction with the TCR with minimal variability compared to the other proteins. It has been shown to be a stable protein to work with in terms of protein-protein interactions. MSP10 is also a potential serological marker and a potential vaccine candidate due to the strong induction of the immune response [42]. The outcomes of this study corroborate the utilization of MSP10 in future malaria vaccine research.

It is interesting to note that hydrophobic interactions usually prevail in protein-protein interfaces. The structural analysis indicates that hydrophobic contacts are commonly used as binding interfaces. As an example, in a T-cell receptor-peptide-HLA complex, all the 26 contacts between a Fab and a neo-antigenic peptide were hydrophobic [58]. This is similar to what we have observed whereby clusters rich on hydrophobic contacts performed well. This is attributed to the fact that hydrophobic residues in the interface reduce solvent exposure and hence a major contribution to binding free energy [58].

Protein to protein binding affinity has been known to be enhanced by hydrophobic interactions. From our findings, it has been established that almost all peptide contacts are hydrophobic in antibody-peptide complexes [58]. Poses in the burying of hydrophobic surfaces in docking are frequently energetically favourable in that, desolvation of nonpolar surfaces gives a large entropic gain. Therefore, clusters that have high hydrophobic interfaces are likely to dominate the scoring. Practically, numerous docking algorithms (such as ClusPro) will give clusters with a high number of hydrophobic contacts as the most populated or lowest-energy. The most populated clusters are likely to represent those poses that have the largest area of hydrophobic interfaces, which is in agreement with the literature. This gives a biological explanation as to why the clusters which “hydrophobic-favored” appeared in our docking.

The molecular docking also provided hints on the stability of the complex structures of antigen-MHC-TCR. The Ramachandran plot also established the stability of the complex structures predicted, with a very high percentage of residues being in the most favored and also allowed regions. There were, however, no more than 90% of the residues being in the most favored regions for the individual models, but the low percentage of disallowed residues (range of 0.3-0.6%) suggests that the complex structures predicted are very well-refined and ready for other methods of analysis. The percentage of this structural validation ensures accuracy of computational predictions for vaccine design. The study of interaction identified stabilizing of the complex structures with major contributions from non-bonded contacts and hydrogen bonds, with the contribution of salt bridges being very prominent in PfCyRPA and PfMSP10. The observation conforms with earlier studies suggesting that hydrophobic and electrostatic interactions are very prominent in binding of antigens with TCR [46].

### Limitations of Docking-based Epitope Prediction

The prediction of epitopes using docking has drawbacks. It should be mentioned that docking full-length antigens to MHC (or TLR) ignores important biological processes. To begin with, peptide cleavage is not performed. Generation of MHC-I epitopes *in vivo* is through the cleavage of proteins by proteasomes and their transportation by TAP [59]. The correct length and termini of peptides only come in the MHC groove. Proteasome cleavage and TAP efficiency prediction methods such as NetCTL contain such tools due to the reason that it isolates which peptides are actually presented [59]. Our simulation ignored proteasomal processing, which may generate peptides that do not form in cells. As a matter of fact, the peptides have to be cut by proteasome and then delivered by TAP followed by MHC binding.

Second, the question of full-length and peptide docking might emerge. T-cells do not recognize full length proteins [60], but only short peptides. According to Tong et al. [60], T cells perceive antigens in form of short peptide fragments in combination with MHC[6]. The docking of complete proteins to MHC is an approximate calculation, since the actual geometry is reliant upon a processed peptide bound to a part of the MHC groove. It is interesting to mention that our docking does not replicate exactly the peptide - MHC interaction.

Third, no refinement of molecular dynamics (MD) was performed. Docking is stiff and results in a static snapshot. These complexes are frequently checked and refined with the help of MD simulations. To illustrate, in a study where a vaccine was docked to TLR4, researchers observed that following 50-100ns MD, the RMSD of the complex had stabilized, indicating that the interface is stable [61]. We did not refined docked complexes using MD; where studies have demonstrated that MD can uncover fluctuation or stabilize the interface.

Finally, there was no HLA coverage analysis of the population. The efficacy of a vaccine is based on the corresponding common HLA alleles. Contemporary vaccine designs actively apply population coverage technologies (e.g. IEDB) to make sure HLA is well represented. Using the IEDB population coverage tool, it has been demonstrated that some specific epitopes may have over 75 percent coverage of some local populations [62]. In our study, HLA allele coverage which is significant context of immunogenicity was not analyzed. We cannot guess what proportion of the population our epitopes may guard without the analysis of coverage. As a matter of fact, peptide vaccine research comes out clearly to state the coverage on a global scale.

## Conclusion

This study highlights the usefulness of computational docking in identifying *P. falciparum* antigens of excellent immunogenic potential as vaccine candidates. Among the complexes studied, PfCSP, PfRipr, and PfMSP10 showed stable and favorable interactions with human MHC–TCR. This is in agreement with their recognized or emerging profile as successful vaccine candidates. Importantly, the MHC-PfSEA-1-TCR complex also scored the lowest energy and it suggests a highly favorable binding conformation. However, its comparatively smaller population of cluster shows that the interaction may be less structurally stable and therefore may be the reason it did not emerge as a significant candidate. Nevertheless, the findings opens the way for further studies on this antigen and in combination with other antigens. In addition, the findings point to the necessity of including multiple antigens of complementary function at different parasite life-cycle stages to deal with problems of variability and immune evasion. The results broadly support the rationale for the development of next-generation multi-epitope malaria vaccines, in which antigen selection needs to be guided by binding affinity as well as conformational stability. Despite the existing experimental validation, further targeted testing is still needed for the predicted complexes, particularly PfSEA-1 and PfMSP10, for immunogenicity and protective efficacy. Through the combination of computational prediction and experimental validation, this approach can accelerate the development of more effective and broadly protective malaria vaccines.

